## Supplementary figures and tables for "Modelling conformational state dynamics and its role on infection for SARS-CoV-2 Spike protein variants"

### Supplementary Material

| | |
|---|---|
| L5F | Mutation not performed – out of range |
| L5F, G476S | Mutation not performed – out of range |
| L5F, D614G | Mutation not performed – out of range |
| L5F, D614G, D839Y | Mutation not performed – out of range |
| L5X | Mutation not performed – out of range |
| L5X, D614G | Mutation not performed – out of range |
| L8V | Mutation not performed – out of range |
| L8V, P1263L | Mutation not performed – out of range |
| L8W, D614G | Mutation not performed – out of range |
| L8X, D614G | Mutation not performed – out of range |

| | |
|---|---|
| H49Q | Mutation performed |
| H49X, D614X | Mutation not performed – nonsense mutation |
| H49X, D614G | Mutation not performed – nonsense mutation |
| H49Y | Mutation performed |
| H49Y, D614G | Mutation performed |
| Y145H, D614G | Mutation performed |
| Q239H, D614G | Mutation performed |
| Q239K, D614G | Mutation performed |
| Q239R, D614G | Mutation performed |
| Q239X, D614G | Mutation not performed – nonsense mutation |
| V367F | Mutation performed |
| V367X | Mutation not performed – nonsense mutation |
| V367F, D614G | Mutation performed |
| G476S | Mutation performed |
| G476S, D614G | Mutation performed |
| V483A | Mutation performed |
| V483F, D614G | Mutation performed |
| V483I | Mutation performed |
| V483X, D614G | Mutation not performed – nonsense mutation |
| D614G | Mutation performed |
| A831S | Mutation performed |
| D614G, A831V | Mutation performed |
| D614G, D839E | Mutation performed |
| D839N | Mutation performed |
| D839X | Mutation not performed – nonsense mutation |
| D614X, D839X | Mutation not performed – nonsense mutation |
| D614G, D839Y | Mutation performed |
| D936H | Mutation performed |
| D936X | Mutation not performed – nonsense mutation |
| D936Y | Mutation performed |
| D614G, P1263L | Mutation not performed – out of range |
| P1263L | Mutation not performed – out of range |
| D614G, P1263X | Mutation not performed – out of range |

**Table S1.** Amino acid mutations associated to 13741 sequences of the Spike protein available on May 08 in COVID-19 Viral Genome Analysis Pipeline, enabled by data from GISAID.

| | |
|---|---|
| Random 1 | L1145F, A1080F, H1088Q |
| Random 2 | I402A |
| Random 3 | F906N, G1035V |
| Random 4 | L865E, N234Y |
| Random 5 | G103H, T881L |
| Random 6 | N74D, L242V, S161W, L335M |
| Random 7 | R1019K, Y636F, L611A, G889I |
| Random 8 | D820R, V213N |
| Random 9 | G971H, G683I, V635P |
| Random 10 | L223M, Q690V, V736C |
| Random 11 | N343C, D290Q, I472P |
| Random 12 | Y741F, S929P |
| Random 13 | F888M |
| Random 14 | N149I, L270S |
| Random 15 | P412H |
| Random 16 | Y365I |
| Random 17 | N17W |
| Random 18 | V1060F, P600L |
| Random 19 | P57Q, V915W, L84W |
| Random 20 | F797A, Q1010W, D1118N |
| Random 21 | T167E |
| Random 22 | Q1005L, A771L |
| Random 23 | T240Y, V656I, F592T, L828K |
| Random 24 | K113Q, Q506H, M697F |
| Random 25 | T599E, E281V, W1102M, N331Y |
| Random 26 | L118C, P330K, F55P |
| Random 27 | I850L, S673R, F1052H, L216P |
| Random 28 | Q414S, P1140Q |
| Random 29 | D737S, W353E, F175R |
| Random 30 | R328G, V512G, E96M, K557H |

**Table S2.** Random mutants accumulating from one to four mutations.

| Variant | $\Delta S_{vib\ (open)} - \Delta S_{vib\ (close)}$ | Predicted Occupancy | | |
| --- | --- | --- | --- | --- |
| | | Open state | Closed State | Difference (Open – Closed) |
| Q14H | 0.345 | 25.836% | 74.164% | -48.328% |
| N61P | 0.324 | 25.835% | 74.165% | -48.330% |
| K417I | 0.467 | 25.806% | 74.194% | -48.389% |
| P491H | 0.374 | 25.804% | 74.196% | -48.391% |
| R355Y | -0.632 | 25.772% | 74.228% | -48.456% |
| Q14F | 0.350 | 25.765% | 74.235% | -48.470% |
| Y369N | 0.379 | 25.731% | 74.269% | -48.538% |
| K417C | 0.500 | 25.725% | 74.275% | -48.549% |
| Q409A | 0.403 | 25.719% | 74.281% | -48.562% |
| F486M | 0.310 | 25.658% | 74.342% | -48.684% |
| R355W | -0.541 | 25.648% | 74.352% | -48.705% |
| I231P | -0.566 | 25.622% | 74.378% | -48.756% |
| G416S | 0.331 | 25.503% | 74.497% | -48.994% |
| K417M | 0.310 | 25.467% | 74.533% | -49.066% |
| G416N | 0.306 | 25.319% | 74.681% | -49.362% |
| P230Y | -0.547 | 25.248% | 74.752% | -49.505% |
| Y489M | 0.422 | 25.168% | 74.832% | -49.663% |
| D111W | -0.547 | 25.162% | 74.838% | -49.677% |
| F464L | -0.575 | 25.030% | 74.970% | -49.941% |
| E465W | 0.361 | 24.977% | 75.023% | -50.046% |
| E465K | 0.322 | 24.896% | 75.104% | -50.208% |
| E465Y | 0.391 | 24.798% | 75.202% | -50.405% |
| D111F | -0.656 | 24.782% | 75.218% | -50.436% |
| Y369V | 0.382 | 24.711% | 75.289% | -50.578% |
| L368N | 0.339 | 24.650% | 75.350% | -50.700% |
| L368G | 0.330 | 24.581% | 75.419% | -50.838% |
| P230D | -0.683 | 24.515% | 75.485% | -50.969% |
| L368S | 0.337 | 24.336% | 75.664% | -51.329% |
| L368C | 0.358 | 24.173% | 75.827% | -51.654% |
| S469W | -0.523 | 23.995% | 76.005% | -52.011% |
| L368A | 0.383 | 23.954% | 76.046% | -52.091% |
| L368P | 0.331 | 23.829% | 76.171% | -52.342% |
| Y369E | 0.501 | 23.744% | 76.256% | -52.513% |
| E465A | 0.360 | 23.694% | 76.306% | -52.612% |
| E465S | 0.336 | 23.051% | 76.949% | -53.897% |

| | | | | |
|---|---|---|---|---|
| G404Y | 0.414 | 22.941% | 77.059% | -54.117% |
| V503D | 0.303 | 22.614% | 77.386% | -54.773% |
| G404H | 0.341 | 22.321% | 77.679% | -55.359% |
| R403N | 0.331 | 21.822% | 78.178% | -56.356% |
| R403P | 0.308 | 21.716% | 78.284% | -56.568% |
| N394K | -0.524 | 21.586% | 78.414% | -56.828% |
| G404N | 0.411 | 21.544% | 78.456% | -56.911% |
| Y421G | 0.455 | 14.552% | 85.448% | -70.896% |
| G232E | -0.707 | 11.578% | 88.422% | -76.844% |
| G232S | -0.686 | 8.996% | 91.004% | -82.008% |
| G232V | -0.713 | 5.508% | 94.492% | -88.983% |
| G232P | -0.647 | 5.042% | 94.958% | -89.917% |
| G232C | -0.664 | 3.065% | 96.935% | -93.870% |
| G232M | -0.609 | 3.033% | 96.967% | -93.935% |
| G232Q | -0.688 | 2.773% | 97.227% | -94.454% |
| G232T | -0.602 | 1.350% | 98.650% | -97.300% |

**Table S3.** Putative mutations and associated $\Delta\Delta S_{vib}$ (in units of $J.K^{-1}$) and predicted occupancies for the open and closed states for the 64 mutants with $\Delta\Delta S_{vib}>0.3$ and the 20 mutants with lowest $\Delta\Delta S_{vib}$ scores.

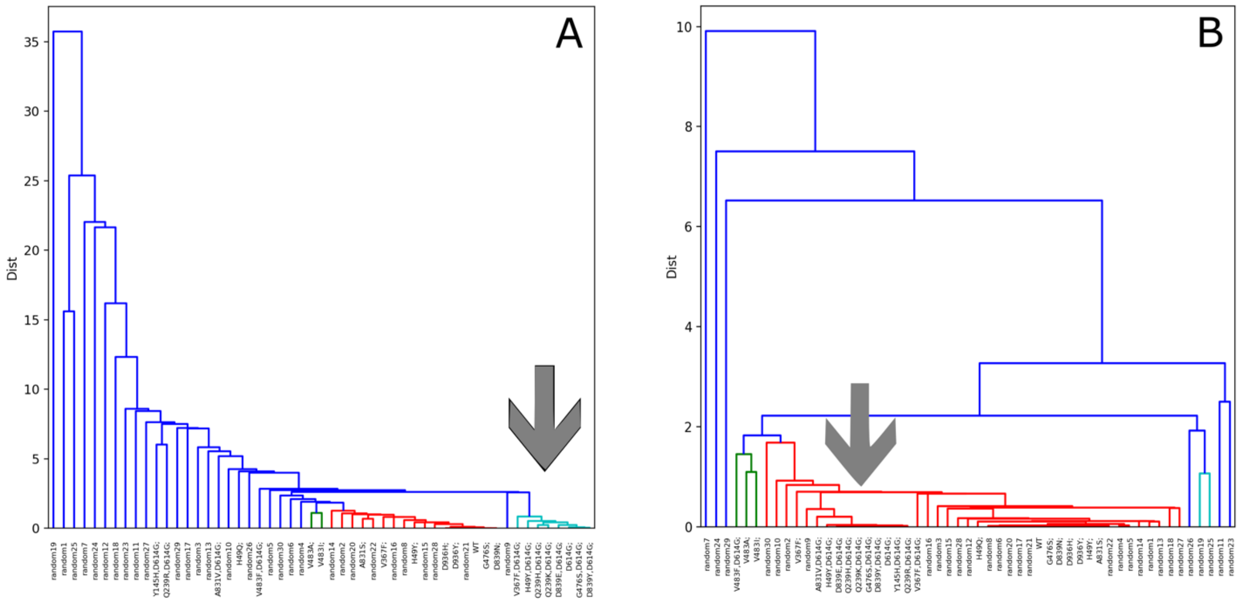


**Figure S1.** Dynamic Signature clustering between WT, 22 mutants observed in nature and 30 random mutants designed, both for the full structure (A) and only for the RBD (B). The grey arrow highlights the clusters containing most of the mutants that have the mutation D614G.

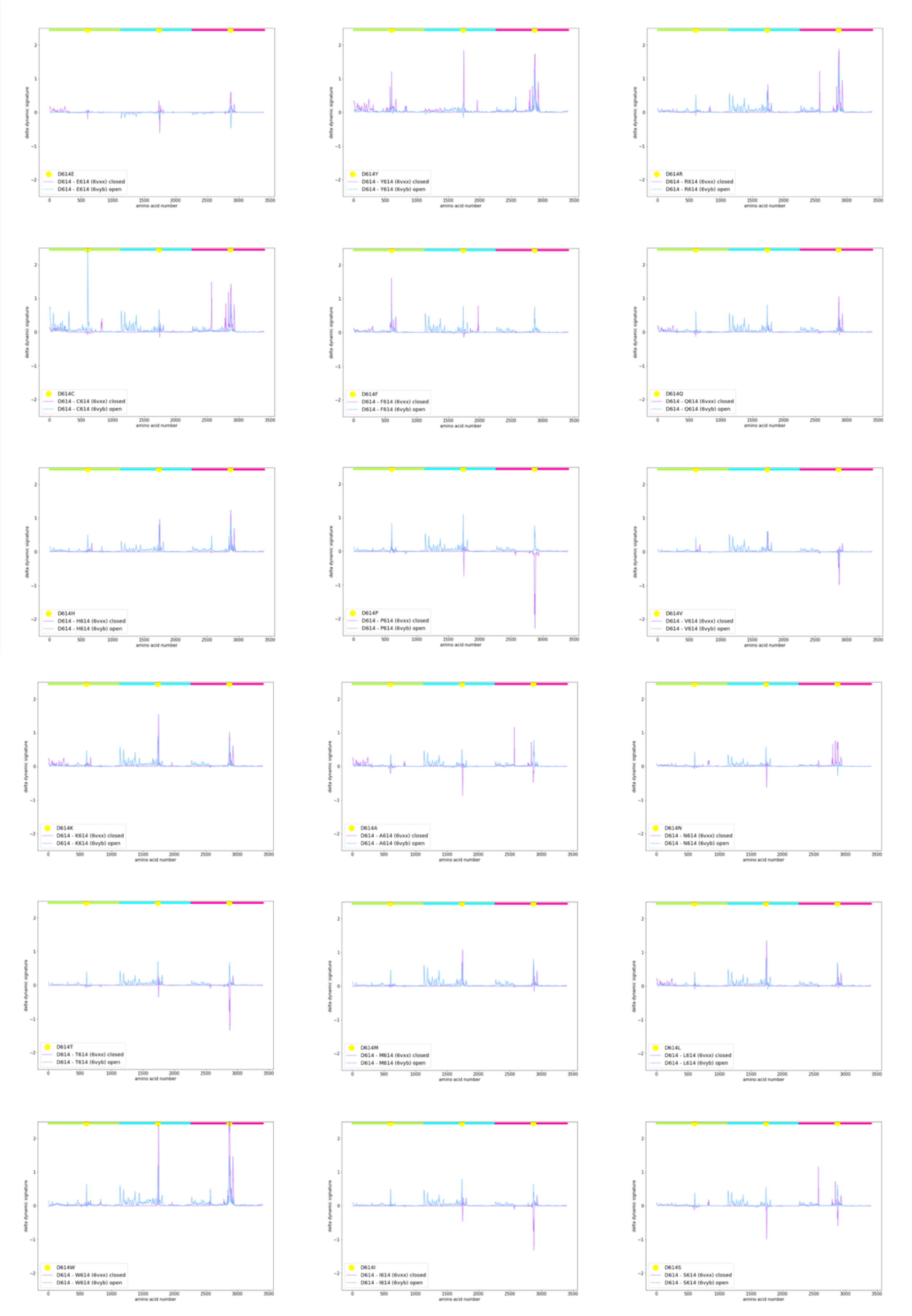


**Figure S2.** Effects on the Dynamic Signatures of the mutation from D614 to each one of the other 18 possible residues.

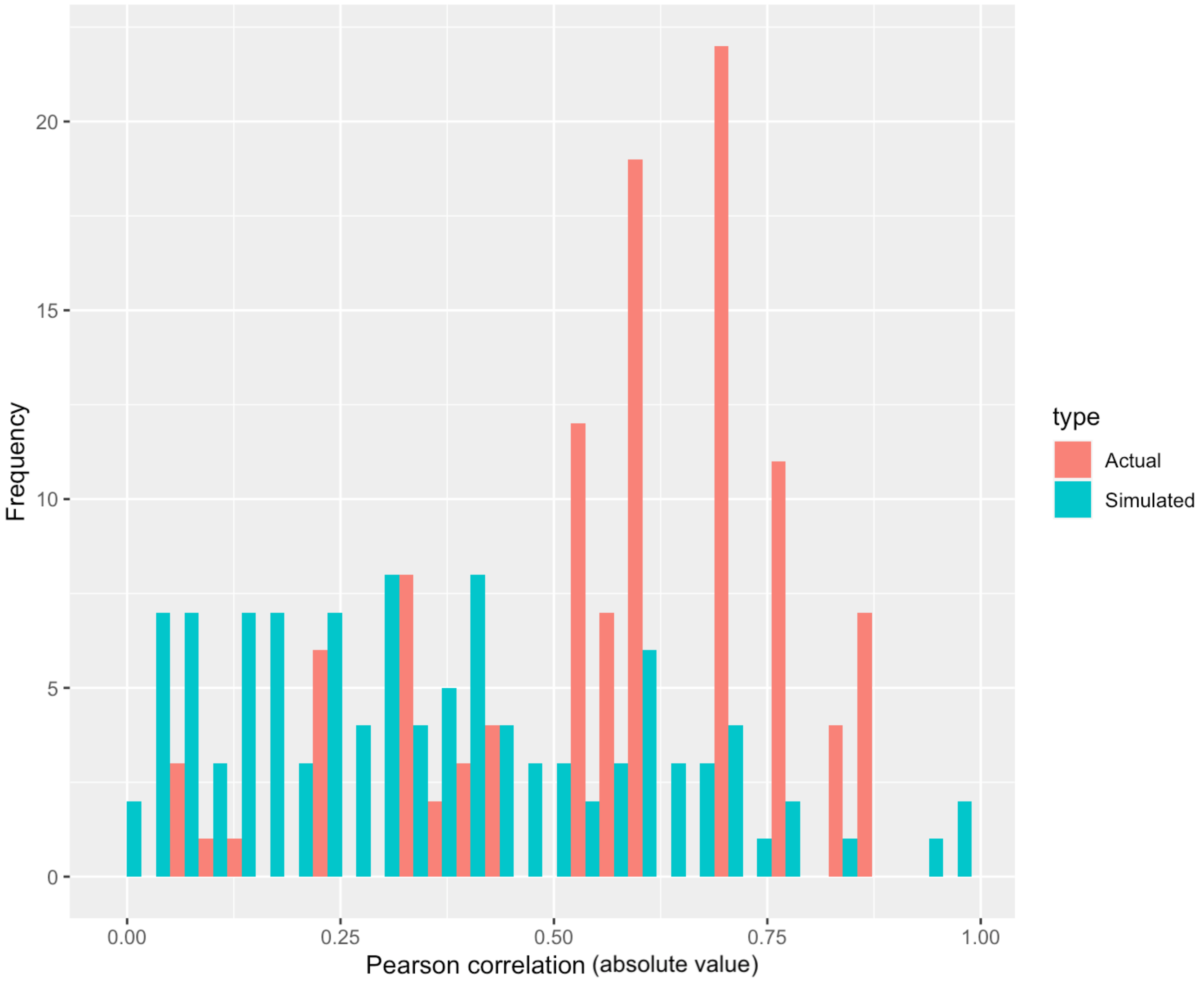


**Figure S3.** Simulated Pearson correlations between gaussian noise vectors of length 6 for the 110 iterations associated to each combination of the parameters k = [0.001, 0.005, 0.01, 0.05, 0.1, 0.5, 1, 5, 10, 50, 100] and $\gamma$ = [0.001, 0.01, 0.1, 1, 10, 100, 1000, 10000, 100000, 1000000].

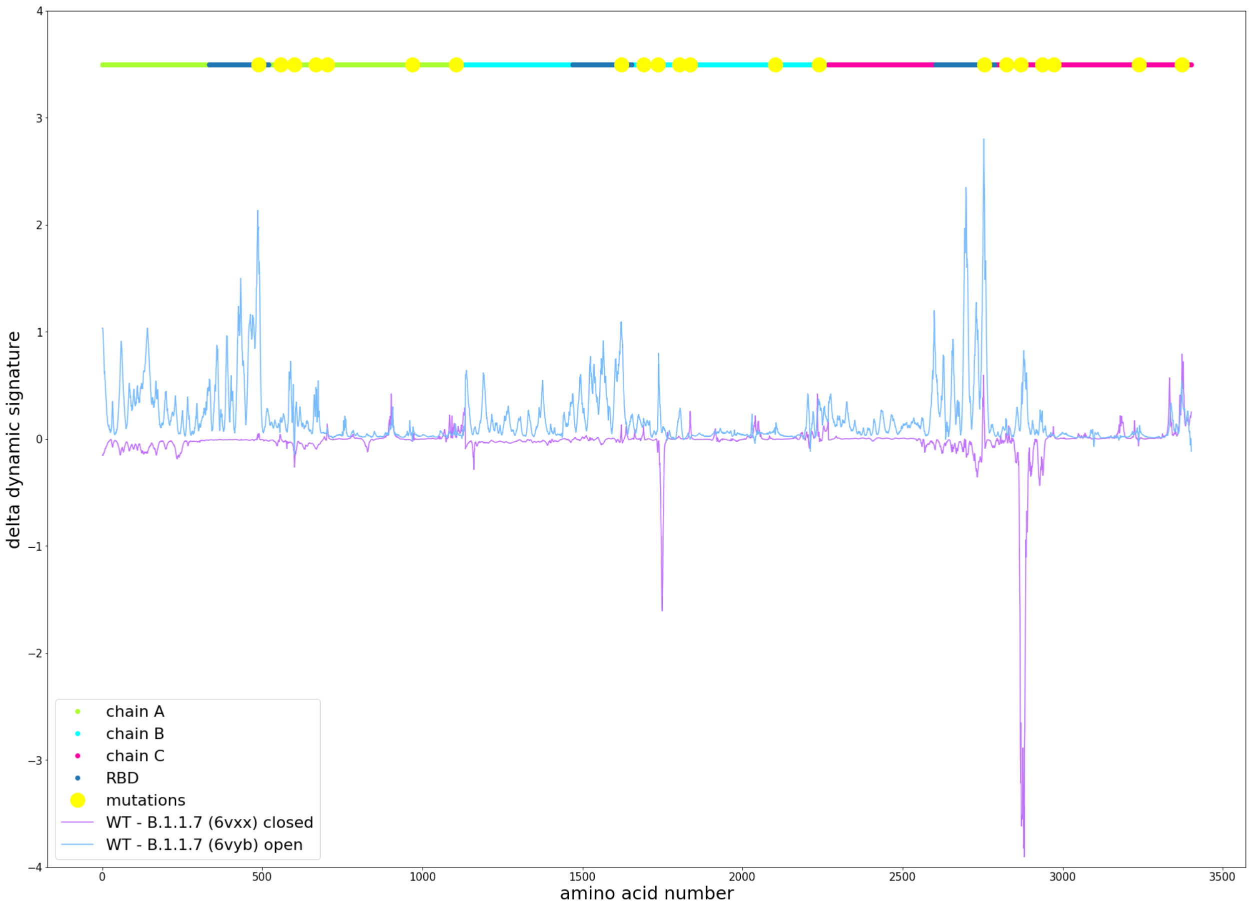


**Figure S4.** Dynamic signature difference for the open (open) and closed (magenta) states relative to the wild type for the B.1.1.7 variant.

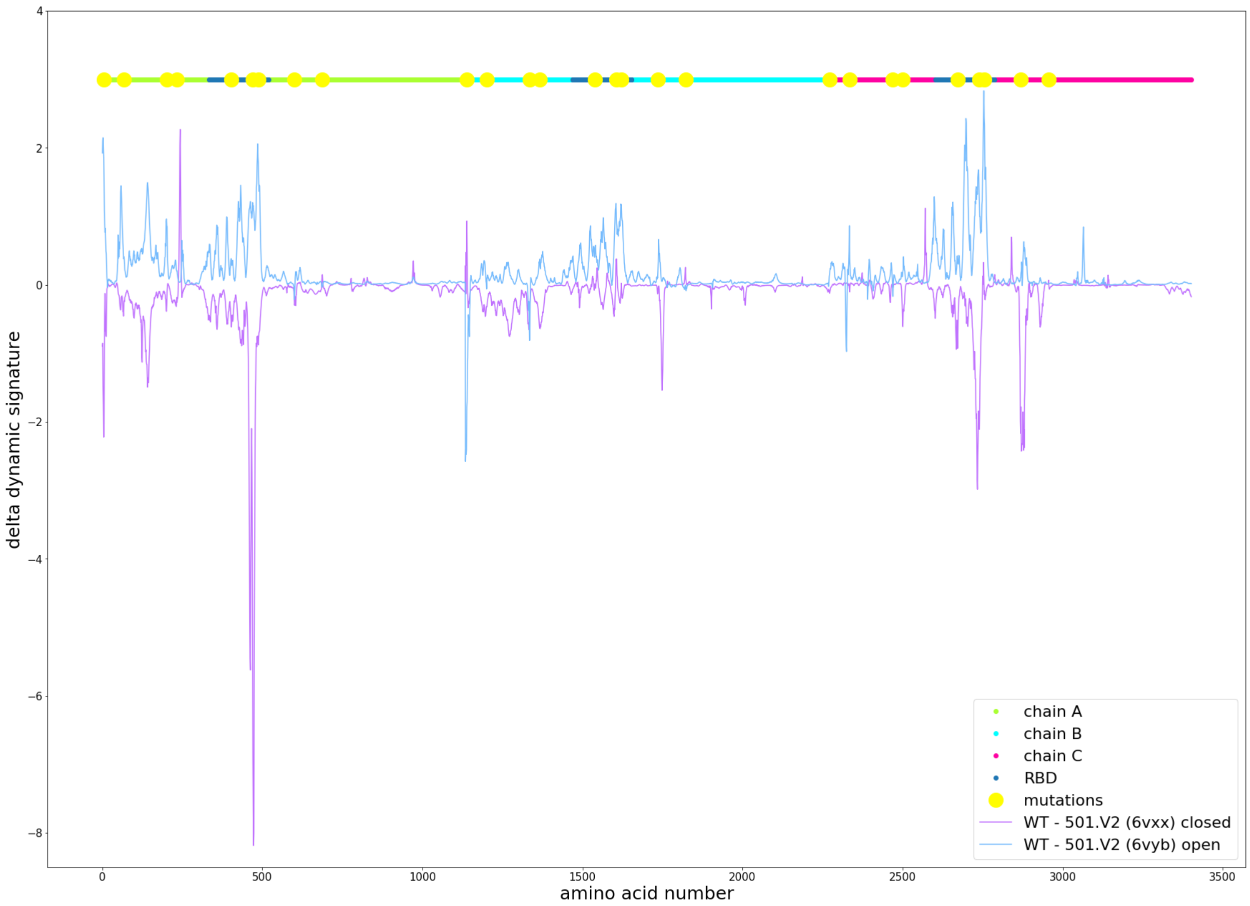


**Figure S5.** Dynamic signature difference for the open (open) and closed (magenta) states relative to the wild type for the 501.V2 variant.
